# Differential expression of deltaFosB in reward processing regions between binge eating prone and resistant female rats

**DOI:** 10.1101/821355

**Authors:** Richard Quansah Amissah, Sandrine Chometton, Juliane Calvez, Genevieve Guèvremont, Elena Timofeeva, Igor Timofeev

## Abstract

Binge eating disorder (BED) is characterized by bingeing and compulsivity. Even though BED is the most prevalent eating disorder, little is known about its pathophysiology. We aimed to identify brain regions and neuron subtypes implicated in the development of binge-like eating in a female rat model. We separated rats into binge eating prone (BEP) and binge eating resistant (BER) phenotypes based on the amount of sucrose they consumed following foot-shock stress. We quantified deltaFosB (ΔFosB) expression to assess chronic neuronal activation during phenotyping. The number of ΔFosB-expressing neurons was: 1) higher in BEP than BER rats in reward processing areas (medial prefrontal cortex (mPFC), nucleus accumbens (Acb), and ventral tegmental area (VTA)); 2) similar in taste processing areas (insular cortex and parabrachial nucleus); 3) higher in the paraventricular nucleus of BEP than BER rats, but not different in the locus coeruleus, which are stress processing structures. To study subtypes of ΔFosB-expressing neurons in the reward system, we performed *in situ* hybridization for glutamate decarboxylase 65 and tyrosine hydroxylase mRNA after ΔFosB immunohistochemistry. In the mPFC and Acb, the proportions of gamma-aminobutyric acidergic (GABAergic) and non-GABAergic ΔFosB-expressing neurons were similar in BER and BEP rats. In the VTA, while the proportion of dopaminergic ΔFosB-expressing neurons was similar in both phenotypes, the proportion of GABAergic ΔFosB-expressing neurons was higher in BER than BEP rats. Our results suggest that reward processing brain regions, particularly the VTA, are important for the development of binge-like eating.

**Significance:** Because ΔFosB expression is associated with a reduction of activity in neurons, a higher expression of ΔFosB in the mPFC, Acb, and VTA of binge eating prone rats compared to binge eating resistant rats suggests a decrease in neuronal activity in these regions, which is consistent with results observed in neuroimaging studies in binge eating disorder patients. This decrease in activity due to ΔFosB expression may underlie the compulsivity and overconsumption of palatable food observed in both our rat model of binge-like eating and binge eating disorder patients.

## Introduction

Eating disorders, namely Anorexia Nervosa (AN), Bulimia Nervosa (BN), and Binge Eating Disorder (BED), cause severe disturbances to eating habits. Hudson, Hiripi (1) reported a lifetime prevalence rate of 0.6% for AN (0.9% of women and 0.3% of men), 1% for BN (1.5% of women and 0.5% of men), and 3% for BED (3.5% of women and 2.0% of men), which suggests that females are more prone to eating disorders than males and that BED is the most prevalent eating disorder [2]. Despite this high prevalence, the pathophysiology of BED is still poorly understood [3].

BED is characterized by eating a large amount of palatable food than would normally be consumed in a discrete amount of time and a loss of sense of control during the bingeing episode [4]. Bingeing episodes are usually triggered by stressful events [5]. While a number of neuroimaging studies, using functional magnetic resonance imaging in humans, showed that BED is associated with an increase in fMRI activity in reward processing brain regions [6–9], others reported a decrease in fMRI activity in similar regions [10–14]. It is therefore unclear whether BED is associated with an increase or a decrease in neuronal activity in these regions.

To study BED, rodent models developed using intermittent access to palatable food and either food restriction or stress, or both, were proposed [15]. Even though each model has its strengths and weaknesses, none of these BED models meets all the criteria defined in the Diagnostic and Statistical Manual of Mental Disorders fifth edition (DSM V). Since BED involves the consumption of palatable food and is triggered by a stressful event, we recently developed a binge-like eating rat model using intermittent access to sucrose solution and foot-shock stress without food restriction, resulting in binge eating prone (BEP; 30% of rats) and binge eating resistant (BER; 30% of rats) rat phenotypes [16].

We hypothesized that reward-, taste-, and stress-mediating brain regions are involved in the development of the binge eating phenotype. The goal of the study was to identify brain regions implicated in the development of binge-like eating, possible differences in neuronal activity between BEP and BER rats in these regions, and the neuron types implicated in these regions. Most studies used c-fos expression to evaluate the effect of acute neuronal stimulation by palatable food consumption in binge-like eating rodents [3, 17]. However, c-fos is transiently expressed and degrades rapidly [18]. We therefore opted for deltaFosB (ΔFosB) because it persists for long periods of time due to its high stability [19] and would last throughout the phenotyping period in our study. ΔFosB accumulates in neurons after chronic stress [20], chronic treatment with drugs [21, 22], and chronic sucrose consumption [23]. Since binge eating involves the consumption of palatable food and is triggered by stress, our aim was to analyze ΔFosB expression in brain regions which process reward (medial prefrontal cortex (mPFC), nucleus accumbens (Acb), and ventral tegmental area (VTA) [24]), taste (parabrachial nucleus (PBN) and insular cortex (IC) [25]), and stress (paraventricular hypothalamic nucleus (PVN) and locus coeruleus (LC) [26]). The results showed that reward processing brain regions are implicated in the development of binge-like eating. We also found that the VTA is the main reward processing region with differential neuronal activation in BEP and BER rats.

## Materials and Methods

All experiments were performed in accordance with the guidelines of the Canadian Council on Animal Care and approved by the Université Laval Committee on Ethics and Animal Research <u>(protocol 2017013)</u>.

### Animals

Forty naïve 45-day-old female Sprague-Dawley rats (body weight: 151-175 g) were purchased from the Canadian Breeding Laboratories (St-Constant QC, Canada) for this study. Each rat was housed individually and maintained on a 12-hour light/dark cycle with the dark cycle starting at 14:00 h in a housing facility with an ambient temperature of 23 ± 1 ºC. Unless otherwise stated, all rats had *ad-libitum* access to tap water and standard rat chow (2018 Teklad Global 185 Protein Diet; 3.1 kcal/g, Harlan Teklad, Montreal, QC). We allowed seven days for acclimatization of rats to the environmental conditions followed by 24-hour access to 10% sucrose solution, one week before the start of experiments, to prevent neophobia to the taste of sucrose solution.

### Generation of binge-like eating rat phenotypes

We generated the binge-like eating rat phenotype as described in a previous study [16]. Briefly, naïve female Sprague-Dawley rats were given one-hour *ad libitum* access to 10% sucrose solution (non-stress session), just at the start of the dark phase, until sucrose consumption was considered stable. This usually requires four to five accesses to 10% sucrose solution. The interval between any two non-stress sessions was at least two days. Afterwards, they underwent three stress sessions which consisted of an unpredictable foot-shock stress immediately followed by one-hour access to sucrose solution. Each foot-shock stress comprised of four foot-shocks with a direct current of 0.6 mA, lasting for 3 s. The equipment for delivering the foot-shock comprises a chamber with a metal grid floor through which electrical current was sent. The inter-shock interval was 15 s. Consecutive stress sessions were separated by at least three days. Since this is a stress-induced binge-like eating model, only consumption of sucrose during stress sessions were used to classify rats as either BEP or BER. For each stress session, using the tertile approach: rats considered high consumers were placed in the upper tertile, average consumers were placed in the middle tertile, and low consumers were placed in the lower tertile. Any rat which appeared at least twice in the upper tertile and never in the lower tertile was considered BEP, while BER rats were rats which appeared at least twice in the lower tertile and never in the upper tertile [16]. In this model, the proportions of rats identified to be BEP, BER, and intermediate (rats considered neither BEP nor BER) are approximately 30%, 30%, and 40%, respectively. In this study, 11 BEP and 12 BER female rats were obtained. They were subsequently divided into two cohorts (n = 6 and 5 for BEP, and n = 6/cohort for BER).

### Test for compulsivity

A modified light/dark box was used to test compulsivity in the first cohort and this test was conducted according to a previously published study [16]. It consists of a dark zone and a light zone. The light zone comprises a 30 cm x 30 cm box made of white Plexiglas while the black zone comprises a 30 cm x 30 cm box made of black Plexiglas. These two zones are connected by a 10-cm wide open door (see Figure 3A). The light zone was brightly illuminated with a light of 300 lx considered aversive to rats [27]. The dark zone was covered with a lid to allow as minimum amount of light as possible to enter (<5 lx). In the light zone, rats had free access to a 10% sucrose solution in a pre-weighed bottle. The experiment was conducted during the dark phase. Rats were first placed in the light compartment facing the spout of the sucrose bottle. The duration of the test session was 10 minutes. In order to distinguish the activity of rats around the sucrose bottle from activity elsewhere in the light zone, a demarcation (14 cm x 8 cm) around the sucrose bottle, called the zone of sucrose, was made. Rats which, despite the obvious aversive light condition, consumed high amounts of sucrose were considered compulsive [28]. The sucrose bottle was weighted before and after the 10 minute-experiment to determine the quantity of consumed sucrose.

### DeltaFosB immunohistochemistry

Three to four days after last access to sucrose solution, rats were anaesthetized using ketamine (160 mg/kg) and xylazine (20 mg/kg). After confirming that rats had no reflex upon pinching, they were intracardially perfused with 100 ml of ice-cold isotonic saline followed by 200 ml of 4% paraformaldehyde (PFA) solution. The rat brains were kept in 4% PFA at 4 ºC for one week. They were then transferred into 20% sucrose in 4% PFA overnight. Using a sliding microtome (Histoslide 2000, Heidelberger, Germany), we cut 30 µm thick coronal sections of brains and kept them at –30 ºC in a sterile cryoprotecting solution made of sodium phosphate buffer (50 mM), ethylene glycol (30%), and glycerol (20%) until they were processed for immunohistochemistry.

The primary antibody used for ΔFosB immunohistochemistry in this study stains both FosB and ΔFosB, but since FosB is known to degrade with time leaving the shorter 37 kD ΔFosB isoform after chronic stimulation [29], we can confidently say that only a minority of the detected staining were contributed by FosB, similar to the antibody used in other studies [22, 30].

Brain sections were first washed in 1% potassium phosphate buffered saline (PPBS) solution followed by treatment with 30% H_2_O_2_ diluted in methanol (1:10). They were washed again in 1% PPBS and blocked for one hour in a solution comprising 0.4% Triton-X, 2% bovine serum albumin, and 1% PPBS. The sections were incubated overnight at 4 °C in the primary rabbit anti-ΔFosB antibody diluted in the blocking solution (sc-48; Santa Cruz Biotechnology, Santa Cruz, CA, 1:1000). The next day, sections were rinsed in 1% PPBS solution, followed by incubation for one hour at room temperature in 1:1500 biotinylated goat anti-rabbit immunoglobulin (Vector Laboratories Inc., Burlingame, CA) diluted in blocking solution. The sections were then rinsed and transferred into a complex of horseradish peroxidase (HRP)-avidin solution (Vector Laboratories Inc., Burlingame, CA) for one hour at room temperature. It was washed with 1% PPBS and then with tris-imidazole solution. To detect staining, a solution containing tris-imidazole, diaminobenzidine (DAB; 0.12 mg/ml), and 0.3% H_2_O_2_ was used. The sections were kept in DAB solution for 10 minutes, rinsed with PPBS, mounted on slides, and cover-slipped with DPX mounting medium.

### Double-labeling for neuron subtypes

To study neurochemical subtypes of neurons which express ΔFosB, we used a glutamic acid decarboxylase 65 (GAD65) probe to identify GABAergic neurons and a tyrosine hydroxylase (TH) probe to identify dopaminergic neurons in brains of the second rat cohort. *In situ* hybridization was performed as described previously [31]. Following ΔFosB immunohistochemistry, sections were mounted on poly L-lysine coated slides and left to dry overnight under vacuum. The sections were subsequently fixed in 4% PFA for 20 minutes, exposed to proteinase K (10 µg/ml in 100 mM Tris-HCl containing 50 mM ethylenediaminetetraacetic acid (EDTA), pH 8.0) for 25 minutes to break down contaminating proteins, acetylated with acetylate anhydride (0.25% in 0.1 M triethanolamine, pH 8.0), and dehydrated by exposure to ethanol solutions of increasing concentration (50, 70, 95, and 100 %). Afterwards, the slides were vacuum dried for at least two hours, followed by addition of 90 µl of the hybridization solution to the slides. This solution contains an antisense ^35^S-labeled cRNA probe against GAD65 or TH. Cover slips were placed on the slides followed by overnight incubation at 55 ºC. After removal of cover slips the following day, the slides were washed in standard saline citrate (SSC; 0.6 M, 60 mM trisodium citrate buffer, pH 7.0), and exposed for 30 minutes to RNase-A at 37 ºC (20 µg/ml in 10 mM Tris-500 mM NaCl containing EDTA). They were then washed in decreasing concentrations of SSC (2X, 10 minutes; 1X, 5 minutes; 0.5X, 10 minutes; 0.1X, 30 minutes at 60 ºC), followed by dehydration in graded concentrations of ethanol. After vacuum drying for two hours, the slides were defatted in xylene and later dipped in NTB2 nuclear emulsion. The slides were exposed for seven days and then developed in D19 developer for 3.5 minutes at 14-15 ºC. They were later fixed in rapid fixer (Eastman Kodak, Rochester, NY, USA) for five minutes. The slides were then washed for one hour under running water, followed by counterstaining with Thionin (0.25%) and dehydration in graded concentrations of ethanol. They were cleared in xylene and cover-slipped following application of DPX mounting medium.

### Quantification of immunoreactive cells

In order to estimate the number of ΔFosB-positive cells in the various regions of interest, we used the Image-Pro Plus Software version 10.0 (Media Cybernetics, Silver Spring, USA). By comparing each brain section with the corresponding section in the Paxinos rat brain atlas [32], the outlines of regions of interest which are relatively small in size (PVN, VTA, LC, and PBN) were made under the 20x objective of the Olympus BX61 microscope (Olympus Canada, Richmond Hill, ON Canada). For these regions, we analyzed the actual brain sections. For regions of interest which are relatively large in size (mPFC, Acb, and IC), sections were first scanned using the TISSUEScope 4000 scanner (Huron Digital Pathology, St. Jacobs, ON, Canada) to obtain high-quality images of sections for subsequent analysis. ΔFosB-expressing neuron quantification was performed automatically. To do this, the Image-Pro Plus software was used to identify objects within the regions of interest. Subsequently, the software was fine-tuned continuously by the experimenter until majority of the objects considered to be neurons in the region of interest were identified by the software. The parameters used were color, area (in pixels: 90–1500), and size (length: 10–90; width: 5–60). At this point, the value of each parameter was noted and applied to all sections containing regions of interest for analysis. The software was then used to automatically identify all similar objects and the number of objects identified was considered as the number of neurons obtained. To verify the results obtained with the automatic counting, we also performed manual cell counting on some brain sections. The results of both automatic and manual counting were similar. For each brain, the number of neurons identified to express ΔFosB was obtained by averaging the number of ΔFosB-expressing neurons per section in regions of interest in both hemispheres of the brain. The regions of interest were the mPFC (prelimbic (PrL) and infralimbic (IL) cortices; +3.72 mm to +2.72 mm from the bregma), Acb (core and shell; +2.28 mm to +0.96 mm), VTA (−4.80 mm to −5.04 mm), IC (+4.2 mm to +0.12 mm), PBN (medial and lateral parts; −8.88 mm to −9.24 mm), PVN (magnocellular and parvocellular parts; −1.72 mm to −1.92 mm), and LC (−9.60 mm to −9.96 mm).

To quantify double-labeled cells, all sections were scanned using the TISSUEScope 4000 scanner to obtain high-quality images of sections and the regions of interest (an example of a typical GAD/ΔFosB-labeled section is shown in Figure 1A). ΔFosB-expressing cells were then identified in all outlined regions of interest (Figure 1B), as previously described. The Image-Pro Plus software defined the coordinates of all identified ΔFosB-expressing cells using the parameters Center X and Center Y. The coordinates of all cells were then exported into Excel files. Similarly, GAD or TH mRNA expression obtained by *in situ* hybridization, which appears as dark silver grains, were also identified based on specific parameters ((in pixels) area: 1–90; size (length): 1–20; size (width): 1– 20; Figure 1C) and their coordinates were exported into Excel files. By using a custom-written MATLAB script, double-labeled cells were identified when there was an overlap of ΔFosB expression and mRNA expression at the same location as shown in Figure 1D. The least number of dark silver grains required for a cell to be considered as double-labeled was set to 5. In addition to the number of double-labeled cells, cells expressing ΔFosB only were also identified using the MATLAB script. Code is available as extended data.

**Figure 1.**
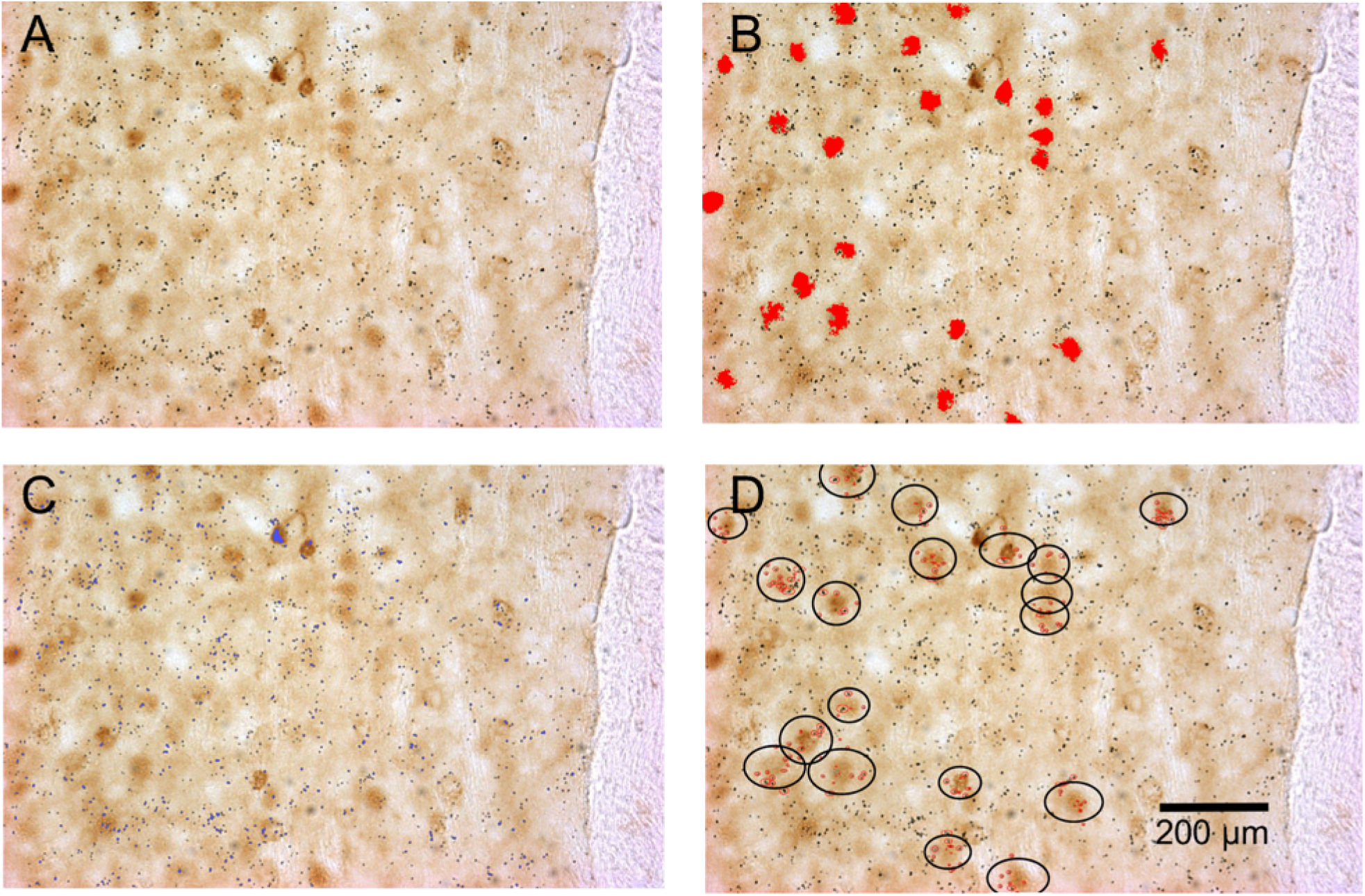
Identification of double-labeled cells (cells expressing both ΔFosB and GAD65 mRNA in this example) using the Image-Pro Plus software and a custom-written MATLAB script. A) An example of a scanned rat brain section showing ΔFosB-expressing cells (dark brown staining) and GAD65 mRNA expression (dark silver grains). B) Identified ΔFosB-expressing cells (red color) using the Image-Pro Plus software. C) Detected GAD65 mRNA expression (red color) using the Image-Pro Plus software. D) Identified double-labeled cells using a custom-written MATLAB script. In the blue circle are ΔFosB-expressing cells that contain more than five dark silver grains (red dots).

### Statistical Analysis

The two-tailed, unpaired student’s t-test was used to compare sucrose intake and time spent in the light zone, dark zone, and zone of sucrose between BEP and BER rats during the 10-minute light/dark box test. Additionally, the two-tailed, unpaired student’s t-test was used to compare the difference in means of the number of ΔFosB-expressing and double-labeled (ΔFosB/GAD65 mRNA and ΔFosB/TH mRNA) neurons in the regions of interest in BEP and BER rats. The ordinary two-way analysis of variance (ANOVA) test followed by the Bonferroni post-hoc test to correct for multiple comparisons was used to compare the total quantities of sucrose solution consumed (dependent variable) by BEP and BER rats (independent variable 1) during sessions with and without foot-shock stress (independent variable 2). The interaction between these two independent variables was also assessed. Data are expressed as mean ± standard error of mean (SEM). Differences in means were considered significant when p < 0.05. The statistical tests were performed using GraphPad version 6.01 (GraphPad Software Inc., La Jolla CA, USA).

## Results

### Sucrose intake during phenotyping

Similar to the sucrose intake of BEP and BER rats generated in the study of Calvez and Timofeeva (16), during one hour sessions, BER rats consistently consumed smaller amounts of 10% sucrose solution compared to BEP rats both during non-stress and stress sessions (Figure 2). BEP rats increased their intake of sucrose solution after foot-shock stress, while the BER rats consumed similar amounts of sucrose solution both during sessions with and without foot-shock stress (p > 0.9999). There was an effect of phenotype (F_1,42_ = 47.41, p < 0.0001) and interaction between phenotype and treatment (F_1,42_ = 5.611, p = 0.0225), but not treatment (F_1,42_ = 3.294, p = 0.0767) on sucrose consumption in BEP and BER rats.

**Figure 2.**
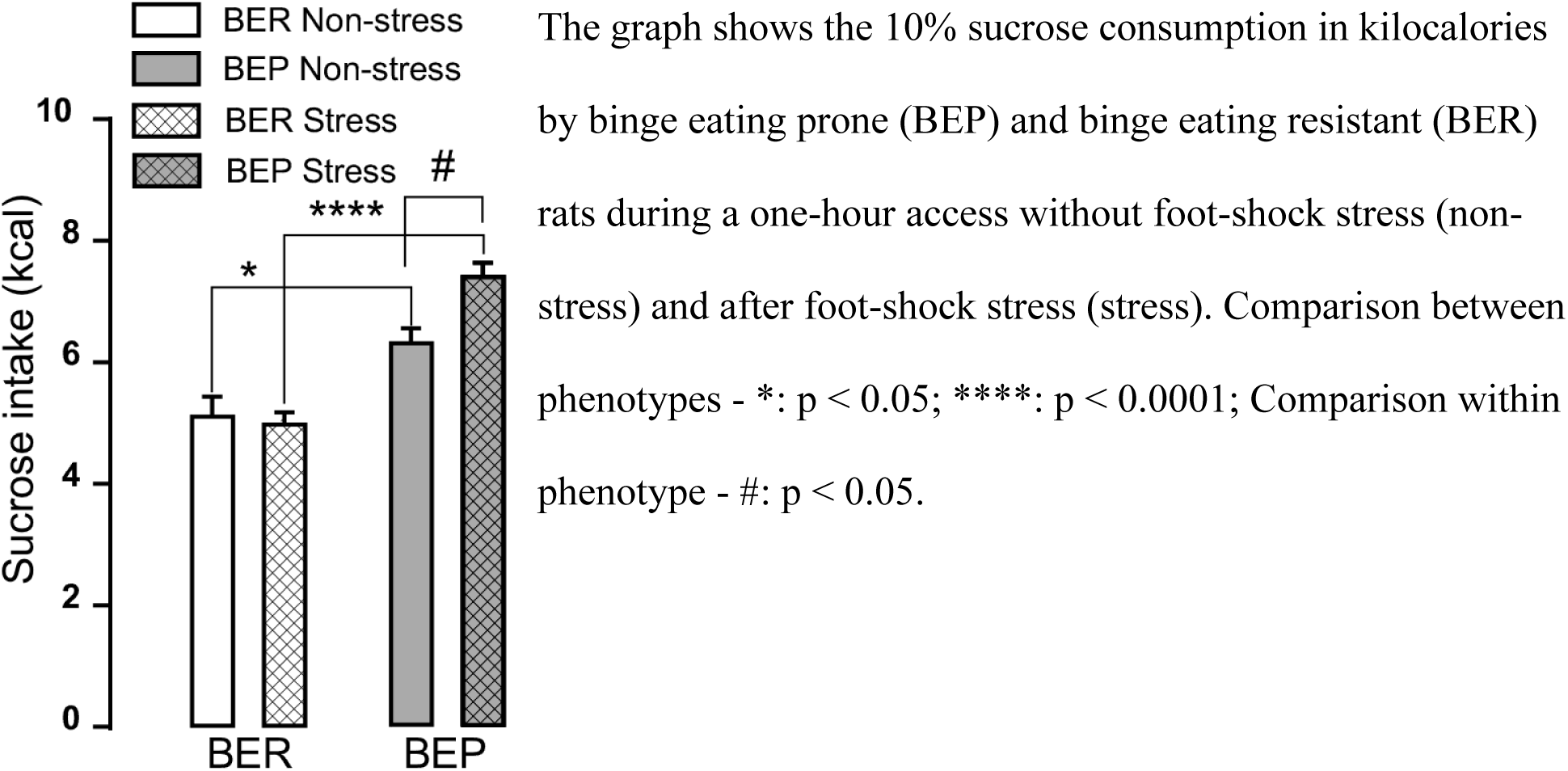
Sucrose intake during phenotyping. The graph shows the 10% sucrose consumption in kilocalories by binge eating prone (BEP) and binge eating resistant (BER) rats during a one-hour access without foot-shock stress (non-stress) and after foot-shock stress (stress). Comparison between phenotypes - *: p < 0.05; ****: p < 0.0001; Comparison within phenotype - #: p < 0.05.

### Compulsive sucrose consumption in BER and BEP rats

We used the light/dark box test (Figure 3A) to assess compulsivity in rats. Rats were allowed to explore the box for 10 minutes with *ad-libitum* access to 10% sucrose solution in the light zone. BEP rats consumed more sucrose than BER rats during the 10-minute *ad-libitum* access to sucrose in the light/dark box (Figure 3B). The zones of interest within the light/dark box were the dark zone, light zone, and zone of sucrose. BER and BEP rats spent similar amounts of time in both the light (Figure 3C) and dark (Figure 3D) zones of the box. However, BEP rats spent significantly more time within the zone of sucrose than BER rats (Figure 3E).

**Figure 3.**
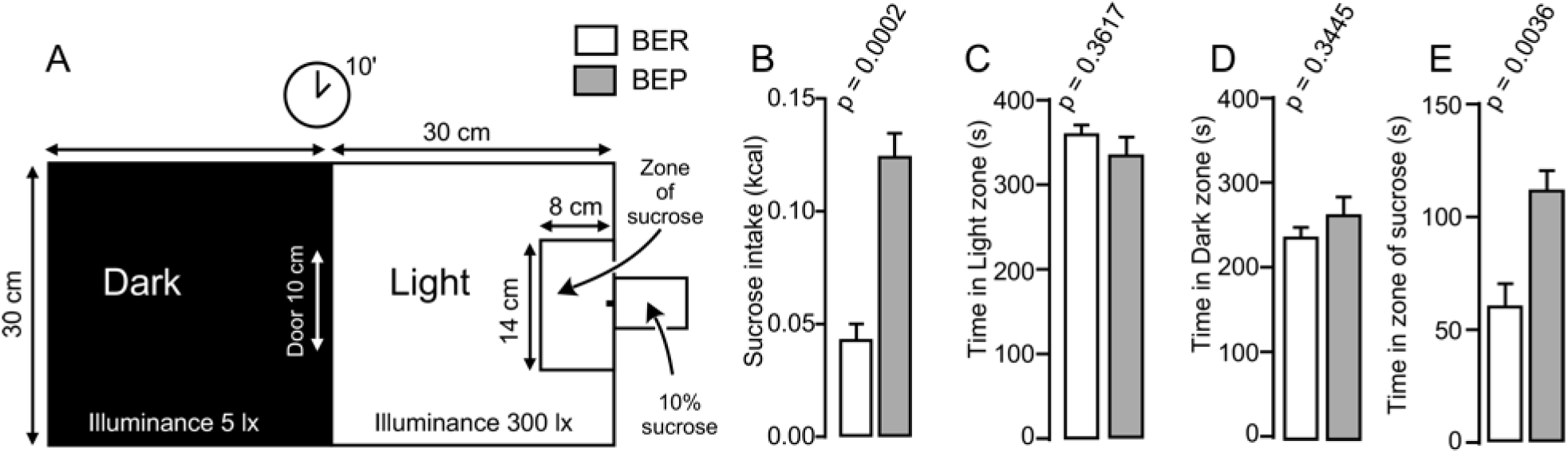
Light/dark box experiment. A) An illustration of the modified light/dark box used during the behavioral experiment. B) Amount of sucrose in kilocalories consumed by binge eating resistant (BER) and binge eating prone (BEP) rats during the 10-minute session in the light/dark box. C) Amount of time in seconds spent by BER and BEP rats in the light zone of the light/dark box. D) Amount of time in seconds spent by BER and BEP rats in the dark zone of the light/dark box. E) Amount of time in seconds spent by BER and BEP rats in the zone of sucrose of the light/dark box. lx: luxes, **: p < 0.01, ***: p < 0.001.

### ΔFosB expression in reward, taste, and stress systems

The number of ΔFosB-expressing cells in all investigated reward processing regions in BEP rats was significantly higher than that in BER rats (Figure 4). A significant difference in the number of ΔFosB-expressing cells was observed in the mPFC, with a higher number of ΔFosB-expressing cells in the PrL and IL of BEP rats compared to BER rats (Figure 4A-D). ΔFosB expression was significantly higher in BEP than in BER rats in the AcbC and AcbSh (Figure 4E-H). There were also more ΔFosB-expressing cells in the VTA of BEP rats compared to BER rats (Figure 4I-K).

**Figure 4.**
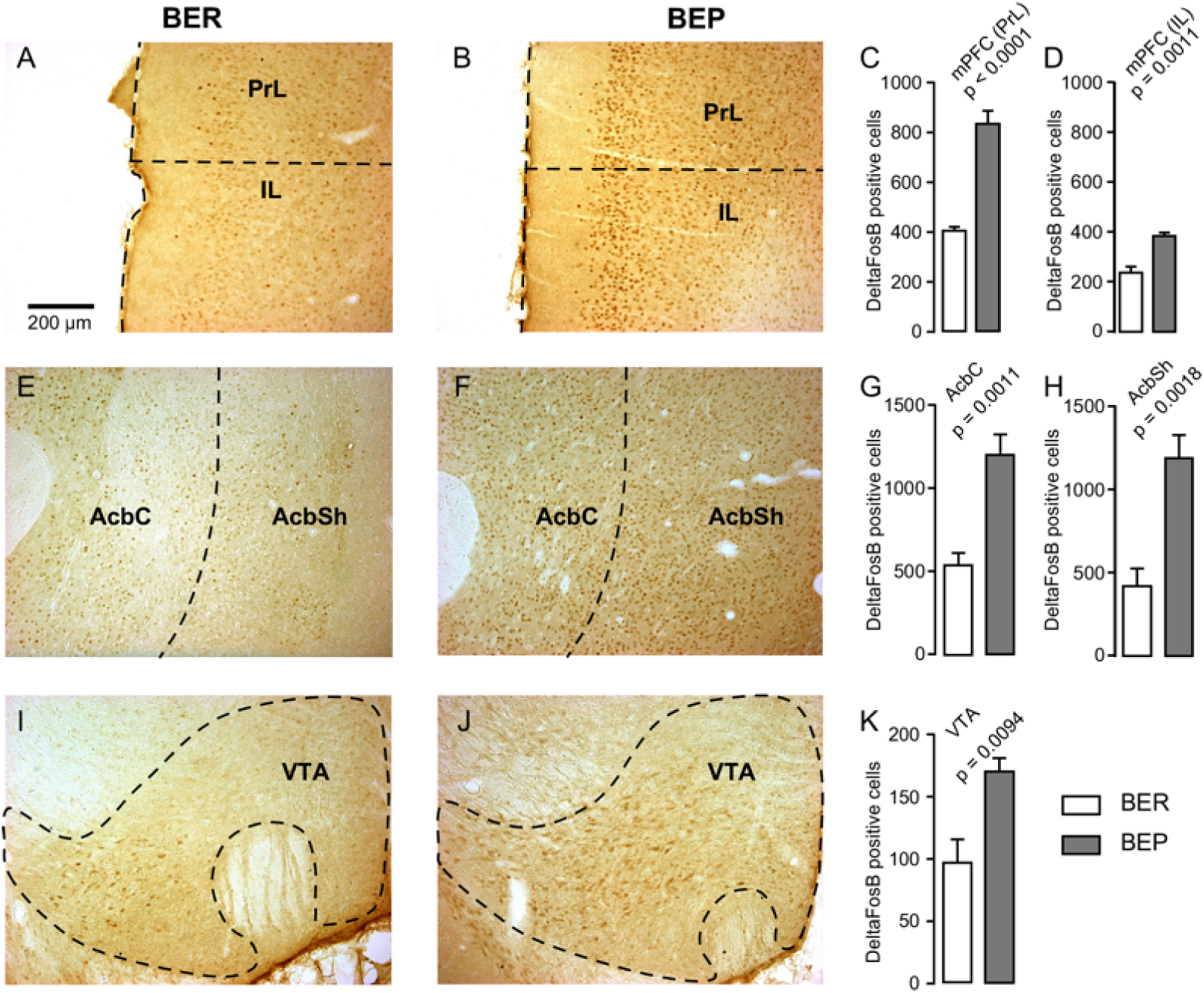
ΔFosB expression in neurons in reward processing regions. A) and B) Images showing ΔFosB-expression in neurons in the prelimbic (PrL) and infralimbic (IL) cortices of the medial prefrontal cortex (mPFC) in binge eating resistant (BER) and binge eating prone (BEP) rats. Inset: A schematic showing the location from which the images in A and B were acquired. C) and D) The number of ΔFosB-positive cells in the PrL and IL of the mPFC in BEP and BER rats. E) and F) Images showing ΔFosB-expression in neurons in the nucleus accumbens core (AcbC) and shell (AcbSh) in BER and BEP rats. G) and H) The number of ΔFosB-positive cells in the AcbC and AcbSh in BEP and BER rats. I) and J) Images showing ΔFosB-expression in neurons in the ventral tegmental area (VTA) in BEP and BER rats. K) The number of ΔFosB-expressing cells in the VTA of BEP and BER rats. **: p < 0.01, ****: p < 0.0001. Scale bar: 200 µm.

ΔFosB expression was also analyzed in a subset of taste processing regions including the IC and PBN. Similar numbers of ΔFosB-expressing cells were identified in both the medial and lateral parts of the PBN in BEP and BER rats (Figure 5A-D), as well as in the IC (Figure 5E-G).

**Figure 5.**
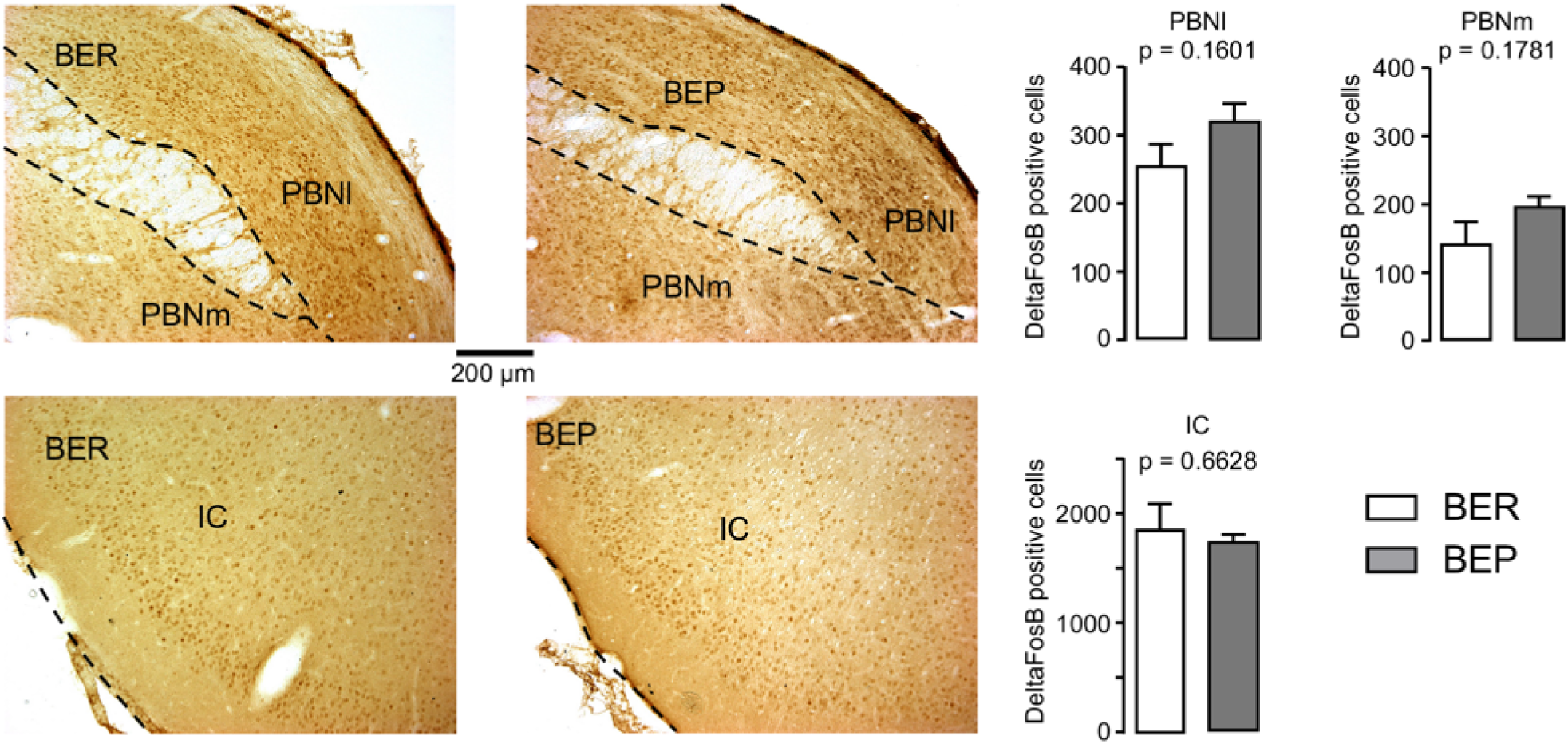
ΔFosB expression in neurons in taste processing regions. A) and B) Images showing ΔFosB-expression in neurons in the lateral (PBNl) and medial (PBNm) parts of the parabrachial nucleus (PBN) of binge eating prone (BEP) and binge eating resistant (BER) rats. C) and D) The number of ΔFosB-positive cells in the PBNl and PBNm in BEP and BER rats. E) and F) Images showing ΔFosB-expression in neurons in the insular cortex (IC) of BEP and BER rats. G) The number of ΔFosB-positive cells in the IC of BEP and BER rats. Scale bar: 200 µm.

We also analyzed ΔFosB expression in two stress processing regions: the LC and PVN (Figure 6). Our analyses revealed that there was a significantly higher number of ΔFosB-expressing cells in both the magnocellular and parvocellular parts of the PVN of BEP rats (Figure 6A-D). However, the expression of ΔFosB in the LC of BEP and BER rats (Figure 6E-G) was similar.

**Figure 6.**
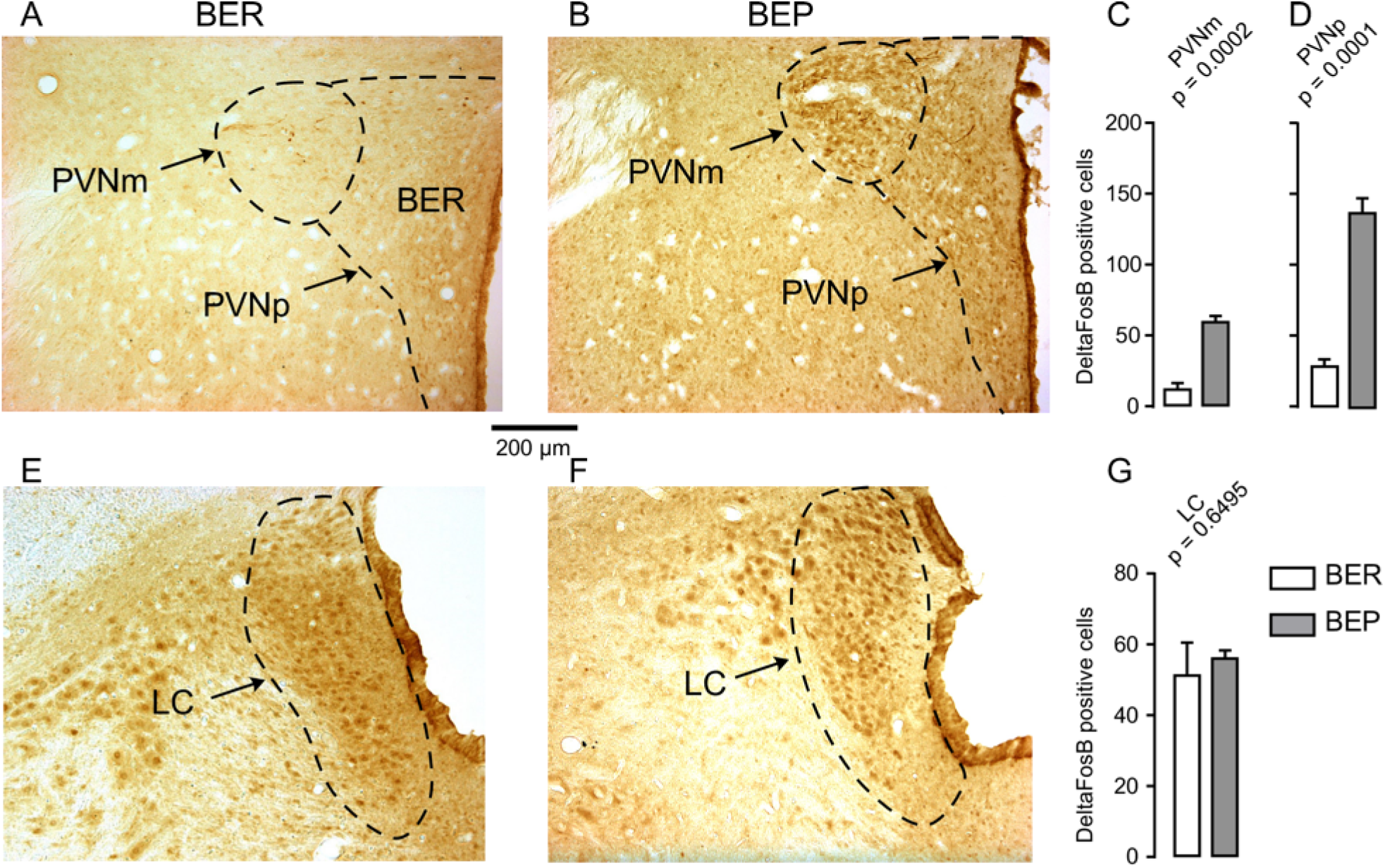
ΔFosB expression in neurons in stress processing regions. A) and B) Images showing ΔFosB-expression in neurons in the magnocellular (PVNm) and parvocellular (PVNp) parts of the paraventricular nucleus of the hypothalamus (PVN) of binge eating prone (BEP) and binge eating resistant (BER) rats. C) and D) The number of ΔFosB-positive cells in the PVNm and PVNp in BEP and BER rats. E) and F) Images showing ΔFosB-expression in neurons in the locus coeruleus (LC) of BEP and BER rats. G) The number of ΔFosB-positive cells in the LC of BEP and BER rats. ***: p < 0.001. Scale bar: 200 µm.

These results show an increase in ΔFosB expression in reward processing areas and in one of the analyzed stress regions, but not in taste processing areas in BEP rats as compared to BER rats.

### Neuronal types implicated in binge-like eating in reward processing regions of the brain

#### Nucleus Accumbens

About 95% of neurons in the Acb are GABAergic cells [33]. Therefore, we investigated whether the increase in ΔFosB expression during our phenotyping was due to the activation of these cells exclusively or other types of neurons. We found that the number of cells expressing both GAD65 mRNA and ΔFosB was higher in the AcbC (Figure 7A) and AcbSh (Figure 7C) in BEP compared to BER rats. For both phenotypes, the proportion of ΔFosB-expressing cells that also expressed GAD65 mRNA was 85-90% in the AcbC (Figure 7B) and AcbSh (Figure 7D), suggesting that ΔFosB expression occurred mainly in GABAergic cells but also in non-GABAergic cells in the Acb of both BEP and BER rats.

**Figure 7.**
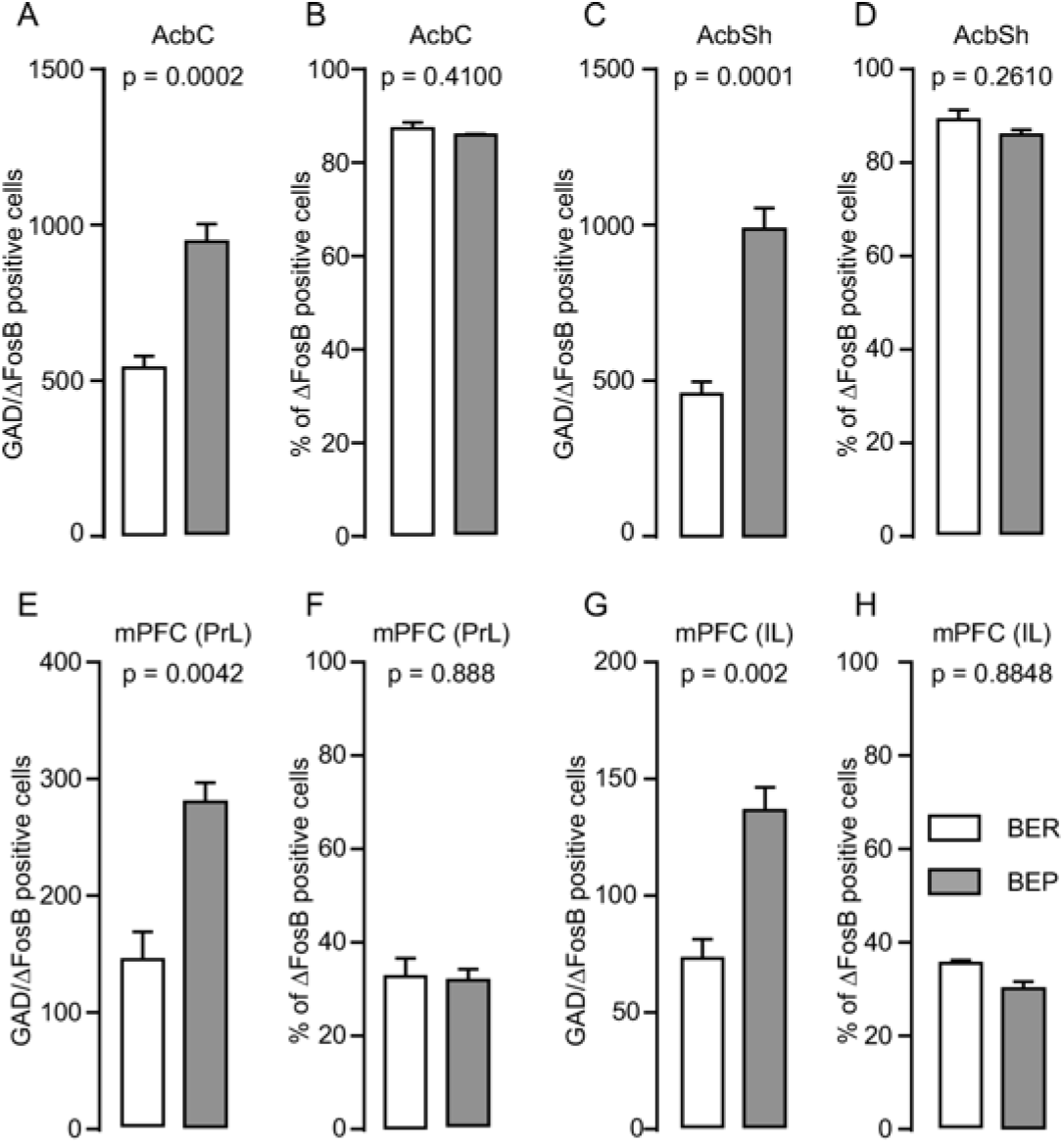
ΔFosB and GAD65 mRNA expression in neurons in the nucleus accumbens (Acb) and medial prefrontal cortex (mPFC) in binge eating prone (BEP) and binge eating resistant (BER) rats. A) and B) The number and percentage of double-labeled cells (ΔFosB cells that express GAD65 mRNA) in the nucleus accumbens core (AcbC) in BEP and BER rats. C) and D) The number and percentage of double-labeled cells in the nucleus accumbens shell (AcbSh) in BEP and BER rats. E) and F) The number and percentage of double-labeled cells in the prelimbic cortex (PrL) of the medial prefrontal cortex (mPFC) in BEP and BER rats. G) and H) The number and percentage of double-labeled cells in the infralimbic cortex (IL) of the mPFC in BEP and BER rats. **: p < 0.01, ***: p < 0.001.

#### Medial Prefrontal Cortex

Gabbott, Dickie (34) reported that about 20% of mPFC neurons are GABAergic, while the remaining are glutamatergic. We found that a higher number of neurons co-expressed ΔFosB and GAD65 mRNA in the PrL of BEP compared to BER rats (Figure 7E). The findings were similar in the IL where more ΔFosB-expressing neurons also expressed GAD65 mRNA in BEP compared to BER rats (Figure 7G). The percentage of ΔFosB-positive neurons that expressed GAD65 mRNA were similar in the PrL (Figure 7F) and IL (Figure 7H) of both BEP and BER rats and was about 30% of the population of the ΔFosB-expressing neurons.

#### Ventral Tegmental Area

Majority of the neurons in the VTA (65%) are dopaminergic neurons, followed by GABAergic neurons which make up 30%, and then glutamatergic neurons which make up about 5% of the total neuron population [35]. In the VTA, ΔFosB-positive cells which also expressed GAD65 mRNA were observed (Figure 8A, B, D, and E). There was no difference in the number of ΔFosB-expressing cells which co-expressed GAD65 mRNA identified in BEP and BER rats (Figure 8C). Interestingly, the percentage of double-labeled cells in the VTA was significantly higher in BER rats compared to BEP rats (Figure 8F).

**Figure 8.**
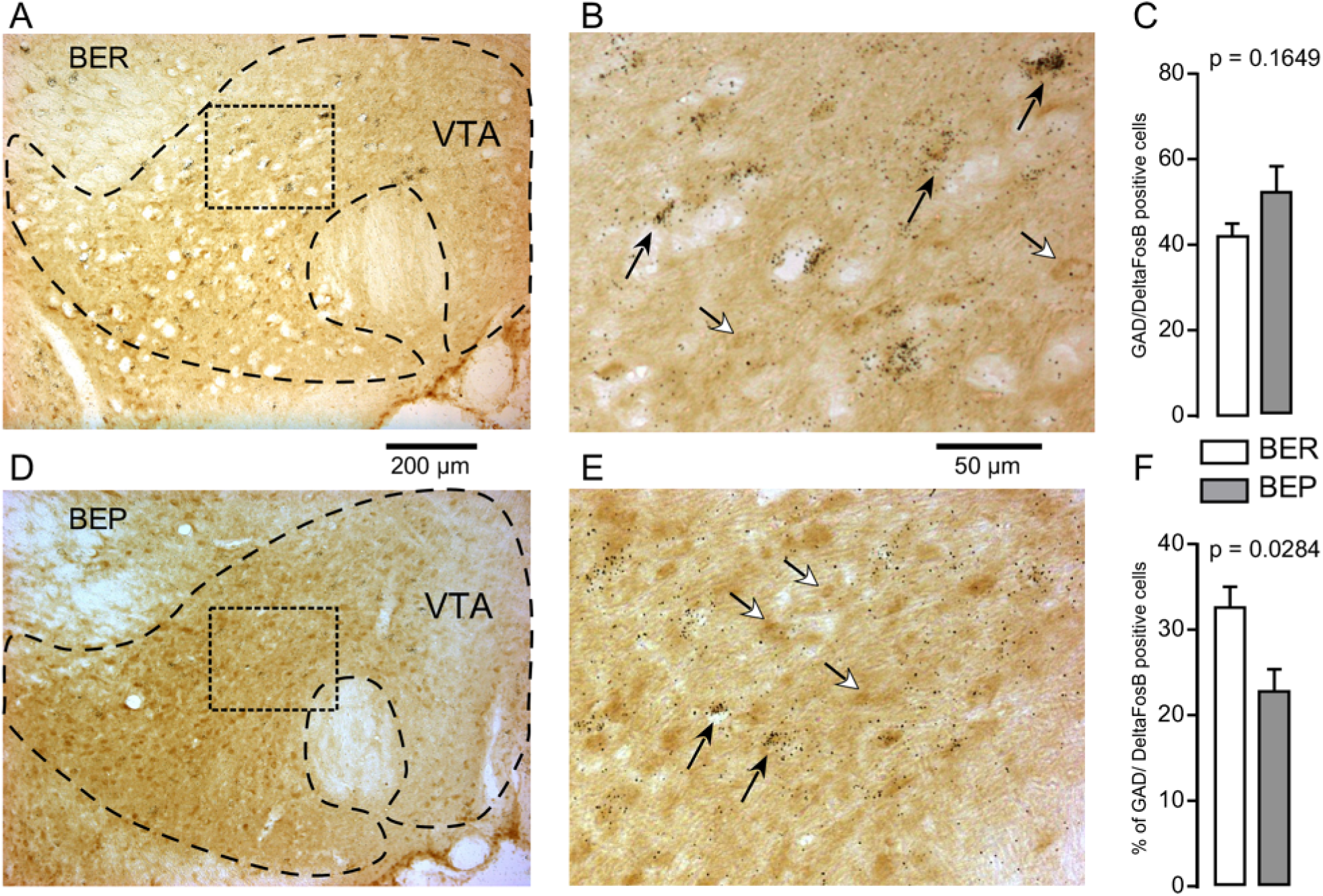
ΔFosB and GAD65 mRNA expression in neurons in the ventral tegmental area (VTA) in binge eating prone (BEP) and binge eating resistant (BER) rats. A) and B) Images showing the labeling of ΔFosB (dark brown) and GAD65 mRNA (dark silver grains) in neurons in the VTA in BER rats. C) The number of ΔFosB/GAD65 mRNA-expressing cells in the VTA in BEP and BER rats. D) and E) Images showing the labeling of ΔFosB and GAD65 mRNA in neurons in the VTA in BEP rats. F) The percentage of double-labeled cells (ΔFosB cells that express GAD65 mRNA) in the VTA of BER and BEP rats. Black arrows point to double labeled neurons, white arrows point to ΔFosB only labelled cells. Scale bar: (A and C) = 200 µm, (B and D) = 50 µm.

Cells which co-expressed ΔFosB and TH-mRNA were observed in the VTA (Figure 9A, B, D, and E). The number of cells which expressed both ΔFosB and TH-mRNA in the VTA of BEP rats was significantly higher than that in BER rats (p = 0.0430; Figure 9C). However, there was no difference in the percentage of ΔFosB-expressing cells which were also positive for TH mRNA (p = 0.2962; Figure 9F) in BEP and BER rats.

**Figure 9.**
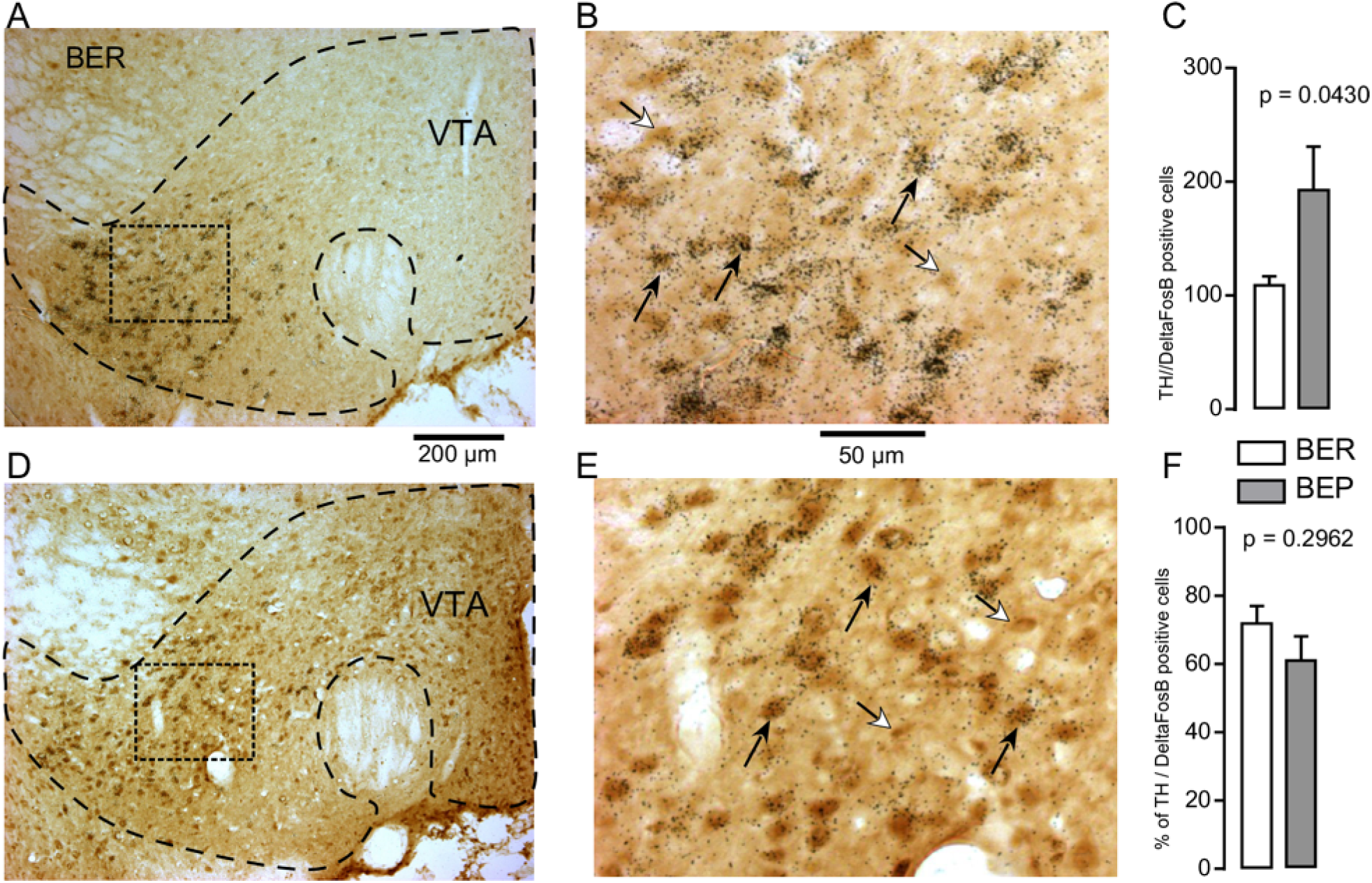
ΔFosB and TH mRNA expression in neurons in the ventral tegmental area (VTA) in binge eating prone (BEP) and binge eating resistant (BER) rats. A) and B) Images showing the labeling of ΔFosB (dark brown) and TH mRNA (dark silver grains) in neurons in the VTA in BER rats. C) The number of ΔFosB/TH mRNA-expressing cells in the VTA in BEP and BER rats. D) and E) Images showing the labeling of ΔFosB and TH mRNA in neurons in the VTA in BEP rats. F) The percentage of double-labeled cells (ΔFosB cells that express TH mRNA) in the VTA of BER and BEP rats. Black arrows point to double labeled neurons, white arrows point to ΔFosB only labelled cells. Scale bar: (A and C) = 200 µm, (B and D) = 50 µm.

## Discussion

BED involves the consumption of a large amount of palatable food and it is usually triggered by stress, which suggests that reward, taste, and stress processing brain regions may be involved [4, 5]. To verify these hypotheses, we evaluated ΔFosB expression in neurons in these regions in our binge-like eating rat model [16]. ΔFosB was expressed after repeated neuronal stimulation, and our binge-like eating rat model was developed using repeated accesses to sucrose and several foot-shock stresses. Our results show that the main brain regions implicated in BED are the reward processing regions (mPFC, Acb, and VTA). In BEP rats, the number of ΔFosB-positive neurons was higher in these regions than in BER rats. Additionally, even though the proportions of non-GABAergic and GABAergic neurons in the mPFC, GABAergic neurons in the Acb, and dopaminergic neurons in the VTA were similar in BEP and BER rats, the proportion of VTA GABAergic neurons involved in binge-eating development was different between the two phenotypes.

ΔFosB expression in the Acb resulted in a reduction in the activity of medium spiny neurons [36]. We observed high ΔFosB expression in the Acb of BEP compared to BER rats. This suggests that there was a significant reduction in the excitability of medium spiny neurons [36] in the Acb of BEP compared to BER rats. In the literature, studies showed that a decrease in neuronal firing in the Acb induced an increase in food consumption whereas stimulation decreased it [37–39]. It was also shown that inhibition of the Acb leads to an increase in the response to [40] and consumption of [23] a reward. Additionally, the brain stimulation of the Acb in mice alleviates binge eating [13]. The high ΔFosB expression in the Acb can be linked with the high sucrose intake observed in our BEP group.

It has been shown that ΔFosB expression in the hippocampus also decreases neuronal activity [41]. We therefore conclude that the activity of ΔFosB-expressing neurons in both the mPFC and VTA would also decrease, and this decrease is greater in BEP than in BER rats. Our results extend the findings of neuroimaging studies which revealed a reduction of activity in the mPFC [11, 12, 14], VTA [42, 43], and Acb [10] in BED patients. Furthermore, a decrease in mPFC activity is associated with compulsivity [44]. BED patients display compulsive behavior, which is associated with a loss of inhibitory control due to hypoactivity in the mPFC [11, 12]. Our light/dark box test showed that BEP rats spent more time in the zone of sucrose and consumed more sucrose than BER rats despite the exposure to the aversive light zone, as was previously shown [16]. It is not surprising that rats consume more palatable food when it is available, but BEP rats consumed more sucrose despite the adverse condition (the light zone) which is abnormal [45]. BEP rats in our study, therefore, exhibit compulsive-like behavior, one of the symptoms of binge eating disorder [4].

Neurons in the taste processing regions (IC and PBN) expressed ΔFosB, and the number of ΔFosB-expressing neurons in these regions was similar in the two phenotypes, which suggests that both phenotypes processed the sucrose taste similarly, even though BEP rats consumed more sucrose than BER rats.

Neurons in the LC, a structure with multiple functions, including stress processing, also expressed ΔFosB. Acute stressful stimuli caused an increase in single unit activity in LC neurons and plasma norepinephrine level [46]. Acute stress activates the LC [47] but as the number of stresses increases, ΔFosB accumulates which reduces the activity of LC neurons. We did not find a difference at the level of LC ΔFosB expression likely because we used repeated foot-shock stresses in our study.

Stress also activates the hypothalamic-pituitary-adrenal (HPA) axis. ΔFosB expression in the PVN was higher in BEP than in BER rats, therefore the activity of these neurons was reduced in BEP rats. We showed previously that these BEP rats displayed a blunted stress-induced activation of the HPA axis, with disruption in the levels of corticosterone and corticotropin-releasing factor (CRF) [48]. This may be due to the high number of ΔFosB-expressing neurons in the PVN of BEP rats observed after repeated stresses.

Our ΔFosB results show that reward processing brain regions are important for developing binge-like eating behavior. Under normal conditions, VTA dopaminergic neurons are activated when a reward is received [49]. These dopaminergic neurons subsequently release dopamine in the mPFC [50] and Acb [51]. The released dopamine activates both glutamatergic and GABAergic neurons in the mPFC [52] and activates GABAergic medium-spiny neurons in the Acb [51]. However, following several stresses (foot-shocks in our experiment), vulnerable rats become binge-eaters [53]. To identify the neuronal subtypes of ΔFosB-expressing neurons in reward processing regions, we performed double-labeling experiments for ΔFosB/GAD65 mRNA to characterize GABAergic neurons and ΔFosB/TH mRNA to characterize VTA dopaminergic neurons. In the Acb, because approximately 95% of neurons are GABAergic medium spiny neurons [33], most of the ΔFosB-expressing neurons were GABAergic in both phenotypes. Since ΔFosB expression reduces neuronal activity [36] and since more ΔFosB/GABAergic neurons were observed in BEP rats than in BER rats, we conclude that the activity of GABAergic neurons in BEP rats was significantly reduced compared to that in BER rats, and this reduction of inhibitory drive can explain their high sucrose consumption [37, 54]. In the mPFC, there were more GABAergic ΔFosB-expressing neurons in BEP rats than in BER rats. However, as the number of neurons which expressed only ΔFosB was also high in BEP rats, the proportion of ΔFosB/GAD65 mRNA-positive neurons was similar in both phenotypes. Since ΔFosB reduces neuronal activity, we conclude that mPFC neuronal activity in BEP rats was significantly reduced compared to that in BER rats, but the proportion of GABAergic and non-GABAergic cells involved are similar for both phenotypes.

Acb medium spiny neurons and both non-GABAergic and GABAergic mPFC neurons express D1 and D2 dopamine receptors and receive projections from VTA dopaminergic neurons [52]. In the VTA, the number of dopaminergic ΔFosB-expressing neurons was higher in BEP rats compared to BER rats, but the proportions of these neurons were similar between the two phenotypes. However, the number of GABAergic ΔFosB-positive neurons was similar in BEP and BER rats. While the expression of ΔFosB in BEP rats was higher, the proportion of GABAergic ΔFosB-expressing neurons was lower in BEP than in BER rats. This difference shows that the VTA is important for the development of binge-like eating. In this region, GABAergic neurons inhibit the activity of dopaminergic neurons. ΔFosB reduces the activity of ΔFosB-expressing neurons [36, 41]. As the proportion of GABAergic ΔFosB-expressing neurons was lower in BEP rats, it suggests that the GABAergic neuronal activity is less reduced in BEP rats compared to BER rats. In other words, the GABAergic neuronal activity in VTA is more present in BEP rats than in BER rats. This implies an overall decrease in the activity of the other neurons in the VTA, and of the other structures that receive VTA GABAergic projections in BEP rats, as previously reported in the VTA of human BED patients [42, 43].

In conclusion, these experiments were designed to analyze, for the first time, ΔFosB expression in different brain regions during the development of binge-like eating in a rat model. We found that the reward system is very important for the development of binge-like eating. In this reward system, the proportions of neuron subtypes involved were similar in the mPFC and Acb, but different in the VTA in BEP and BER rats. The results suggest that these differences in proportion in the VTA may play an important role in bingeing.

## Acknowledgement

This work was funded by the Natural Sciences and Engineering Research Council of Canada (NSERC; E.T., grant 1295926) and the Canadian Institute of Health Research (CIHR; E.T., grant 102659, I.T. grant 136969). We would like to thank Christophe Lenglos for providing the MATLAB scripts for the analyses of deltaFosB, glutamate decarboxylase 65, and tyrosine hydroxylase mRNA expression in this study.

## Extended data

Code for the double labeling detection is uploaded to the journal website.

The code can be run in Matlab. It was used to detect ΔFosB only or double-labelled cells.

